# Nonpolar Residues in the Presumptive Pore-Lining Helix of Mechanosensitive Channel MSL10 Influence Channel Behavior and Confirm a Non-Conducting Function

**DOI:** 10.1101/264283

**Authors:** Grigory Maksaev, Jennette M. Shoots, Simran Ohri, Elizabeth S. Haswell

## Abstract

Mechanosensitive (MS) ion channels provide a universal mechanism for sensing and responding to increased membrane tension. MscS-Like(MSL)10 is a relatively well-studied MS ion channel from *Arabidopsis thaliana* that is implicated in cell death signaling. The relationship between the amino acid sequence of MSL10 and its conductance, gating tension, and opening and closing kinetics remain unstudied. Here we identify several nonpolar residues in the presumptive pore-lining transmembrane helix of MSL10 (TM6) that contribute to these basic channel properties. F553 and I554 are essential for wild type channel conductance and the stability of the open state. G556, a glycine residue located at a predicted kink in TM6, is essential for channel conductance. The increased tension sensitivity of MSL10 compared to close homolog MSL8 may be attributed to F563, but other channel characteristics appear to be dictated by more global differences in structure. Finally, MSL10 F553V and MSL10 G556V provided the necessary tools to establish that MSL10’s ability to trigger cell death is independent of its ion channel function.

## Introduction

The ability to respond to mechanical stimuli is an ancient and intrinsic property of cells ^1,2^. One of the most universal mechanisms for mechanotransduction is the use of mechanosensitive (MS) ion channels. MS channels are oligomeric protein structures embedded in the lipid bilayer, and their primary function is to form a conductive pore in response to increased lateral membrane tension or force transduced from cytoskeletal filaments ^3^. MS channels mediate the perception of external mechanical stimuli (touch, gravity, vibration) and internal mechanical stresses in plants, animals, and bacteria ^4–6^. Mammalian MS channel dysfunctions are associated with numerous pathologies ^7^ and both mammalian and bacterial MS channels are under investigation as drug targets ^8–10^. It is therefore important for both basic and for applied reasons to determine the molecular mechanisms of MS ion channel function, including the relationship between channel structure and its conductance, gating tension, and opening and closing kinetics.

We already have considerable insight into these questions, in part due to decades of research into the structure and function of a MS ion channel from *Escherichia coli*, the Mechanosensitive ion channel of Small conductance (*Ec*MscS). *Ec*MscS is directly opened by membrane tension ^11^, has a unitary conductance of 1.2 nS in giant *E. coli* spheroplasts ^12,13^ and demonstrates a slight preference for anions (1.2-3 fold, summarized in ^14,15^). The primary physiological function of *Ec*MscS is to promote bacterial survival when subjected to hypoosmotic shock ^13,16^. *Ec*MscS also shows inactivation behavior whereby sustained tension leads to a non-conductive state of the channel that cannot be opened again until a period of recovery ^17–21^.

A wealth of structural information on MscS is available, derived from several members of the MscS family from different bacterial species. Crystal structures thought to represent the conducting state of *Ec*MscS or the non-conducting states of MscS from *E. coli, Thermoanaerobacter tengcongensis*, and *Helicobacter pylori* ^22–24^ have been determined. A cryoelectron micrograph structure of MscS homolog from *E. coli*, YnaI, has also been reported ^25^. These structures reveal that MscS forms a homoheptamer with a transmembrane (TM) domain localized to the inner *E. coli* membrane and a cytoplasmic “vestibule”. Each subunit contains an N-terminal domain comprised of three TM helices, and a soluble C-terminal domain. The most C-terminal of the TM helices, TM3, lines the permeation pore. It comprises two regions, TM3a and TM3b, which are separated by a distinctive kink at residue G113. Other key residues include L105 and L109, which form the narrowest constriction of the closed or non-conducting pore, and G121, which is thought to be critical to closed state formation^2217^.

A comparison of the open state versus closed state structures suggest that gating involves swinging a tension-sensitive paddle made up of the TM1/TM2 helices and twisting TM3a about G113. This motion allows L105 and L109 to move out of the pore. Mutational analyses support important roles for L105 and L109 ^26,27^ and have shown that G104, A106 and G108 play critical roles in channel gating ^28,29^. A single mutation in this region, A106V, locks the channel in an open conformation, and was used to obtain the first open-state crystal structure of *Ec*MscS^30^. Mutating other residues in TM3, such as S114, L118, A120, L123, F127 ^31^ and Q112, A120 ^16^ has less dramatic impact on channel properties, only modulating its gating and inactivation kinetics. However, changing the kink-forming residue G113 to alanine prevents inactivation and in combination with G121A severely alters channel opening and closing ^17^. Residues F68 and L111 may form a force-transmitting clutch between TM2 and TM3, transmitting membrane tension sensed by the TM1/TM2 paddle to the pore-forming TM3. ^32^ Surprisingly, the channel’s weak ion selectivity is governed by its cytoplasmic cage, rather than the pore-lining TM3 ^33,34^. In summary, a combination of modeling and functional assays now provide a general understanding of the conformational changes that ultimately result in channel gating for *Ec*MscS ^6,35,36^.

These insights into the structural basis of *Ec*MscS mechanosensitivity provide a strong foundation for studying homologs of *Ec*MscS, which are found in all kingdoms of life ^6,35,36^. The domain conserved among all these MscS family members is limited to the pore-lining helix and about 100 amino acids of the following soluble domain. The number of predicted transmembrane domains and the structure of N- and C-termini are highly variable. The channel behavior and physiological function of multiple *Ec*MscS homologs from other bacterial species, archaea, fission yeast, green algae and land plants have been reported ^2,37–40^. These channels are all mechanically gated and generally function in hypoosmotic stress relief, but have a range of conductances, ion channel selectivities and play different physiological and developmental roles.

The ten *Ec*MscS homologs encoded in the genome of the model land plant *Arabidopsis thaliana* have been named MscS-Like or MSL channels ^41–49^. MSL proteins exhibit diverse tissue expression patterns, subcellular localizations and domain structures ^50^. To date, MSL1, MSL8 and MSL10 are the best-characterized MSLs in terms of ion channel physiology. All three provide tension-gated ion channel activities in native plant cells ^5^ and/or when expressed in heterologous systems ^51^. MSL1 is localized to the mitochondrial inner membrane ^43–45^, while MSL8 and MSL10 are primarily localized to the plasma membrane ^45^. The unitary conductances of MSL8 and MSL10 expressed in *Xenopus* oocytes are approximately 60 pS and 105 pS, respectively (compare to 340 pS for *Ec*MscS expressed in oocytes) ^44,51^. MSL8 and MSL10 have a slightly higher preference for anions (P_Cl_: P_Na_ = 5.6–6.3) than *Ec*MscS. Another intrinsic feature of an MS ion channel is its tension sensitivity, defined as the amount of tension applied to the membrane required for channel opening. The tension at which a channel opens may or may not be the same as the tension at which it closes. Different opening and closing tensions lead to an asymmetric gating profile, and this phenomenon is referred to as hysteresis. Both MSL10 and MSL8 have lower tension sensitivity than *Ec*MscS, and both exhibit strong hysteresis, with much higher opening than closing pressures ^43,44,52,53^, while *Ec*MscS does not. We interpret this to mean that, once opened, MSL8 and MSL10 are very stable and do not close until most of the tension is relieved ^43,44^.

In terms of physiological function, MSL8 appears to serve in a role analogous to that of MscS, as it protects pollen from multiple hypoosmotic challenges associated with pollen development and function ^43^. MSL8 is primarily localized to the plasma membrane of pollen grains, where it is required for full survival of rehydration, germination, and tube growth. Two lesions in the presumptive pore-lining helix of MSL8, I711S and F720L, alter channel behavior and fail to complement these mutant phenotypes. These observations link ion flux through the MSL8 channel to protection from osmotic stresses during pollen development ^44^.

Much less clear is the functional role of MSL10. MSL10 is required for the predominant MS ion channel activity in root cells ^53^, but to date no other loss-of-function phenotype has been established. Fortunately, gain-of-function phenotypes have been revealing; overexpression of MSL10 leads to cell death, as does a single ethyl methanesulfonate-induced point mutation in the MSL10 C-terminus, S640L (*rea1*) ^51^. Over-expression of the soluble N-terminus of MSL10 is sufficient to induce cell death in tobacco epidermal cells ^54,55^. Thus, all existing data suggest that MSL10 is a multifunctional MS ion channel that is capable of mediating adaptation to hypoosmotic shock in the short-term (reducing pressure by releasing osmolytes) <u>and</u> of signaling to change cellular state in the long-term (inducing cell death in response to biotic or abiotic stress). However, more structural information about MSL proteins and their pore-forming region is required to fully and directly test the possibility that MSL10 has a non-conducting function.

To gain additional information about the structure of the channel pore, the mechanism of gating, and how ion flux through the channel is related to its genetic functions, we used mutational analysis and single-channel patch-clamp electrophysiology to identify residues in the presumptive pore-lining domain of MSL10 that are important for channel function, including tension sensitivity, conductivity, and stability of the open state. We then tested two tension-insensitive mutants for the ability to induce cell death in a previously established transient expression assay. These data provide critical information about the structural component of tension-sensitive ion transport, and a useful comparison to *Ec*MscS and other MS channels in animals and bacteria. In addition, these mutant MSL10 channels provide tools for studying the relationship between tension sensitivity, open state stability, ion flux, and cell death signaling.

## RESULTS

### Selection of potential pore-disrupting residues

To identify residues likely to be important for MSL10 ion channel activity, we performed a sequence alignment between the region of highest homology between *Ec*MscS, MSL10 and MSL8 (Figure 1A). In the *Ec*MscS crystal structure, this sequence forms transmembrane (TM) helix 3, the domain that lines the channel pore ^54^. *Ec*MscS TM3 is 33 amino acids long and has a pronounced kink at G113 that splits it into TM3a and TM3b. The analogous sequence in MSL10 is its most C-terminal TM helix, TM6. We therefore generated a hypothetical structure of TM6 using the I-TASSER prediction server ^22^ and the *Ec*MscS closed state crystal structure (2OAU:A, ^56^) as a template (Figure 1B). These data, along with a topology prediction from the ARAMEMNON server ^22,57^, support the overall topology for MSL10 shown in Figure 1C. Based on this alignment, we identified four key classes of residues within the TM6 sequence that were likely to contribute to MSL10 channel function: 1) multiple phenylalanine residues, 2) the putative glycine kink, 3) several nonpolar TM6 residues that differ between MSL10 and MSL8, and 4) an isoleucine residue known to play a role in MSL8 function. Those residues selected for study are indicated with circles in Figure 1A.

**Figure 1.**
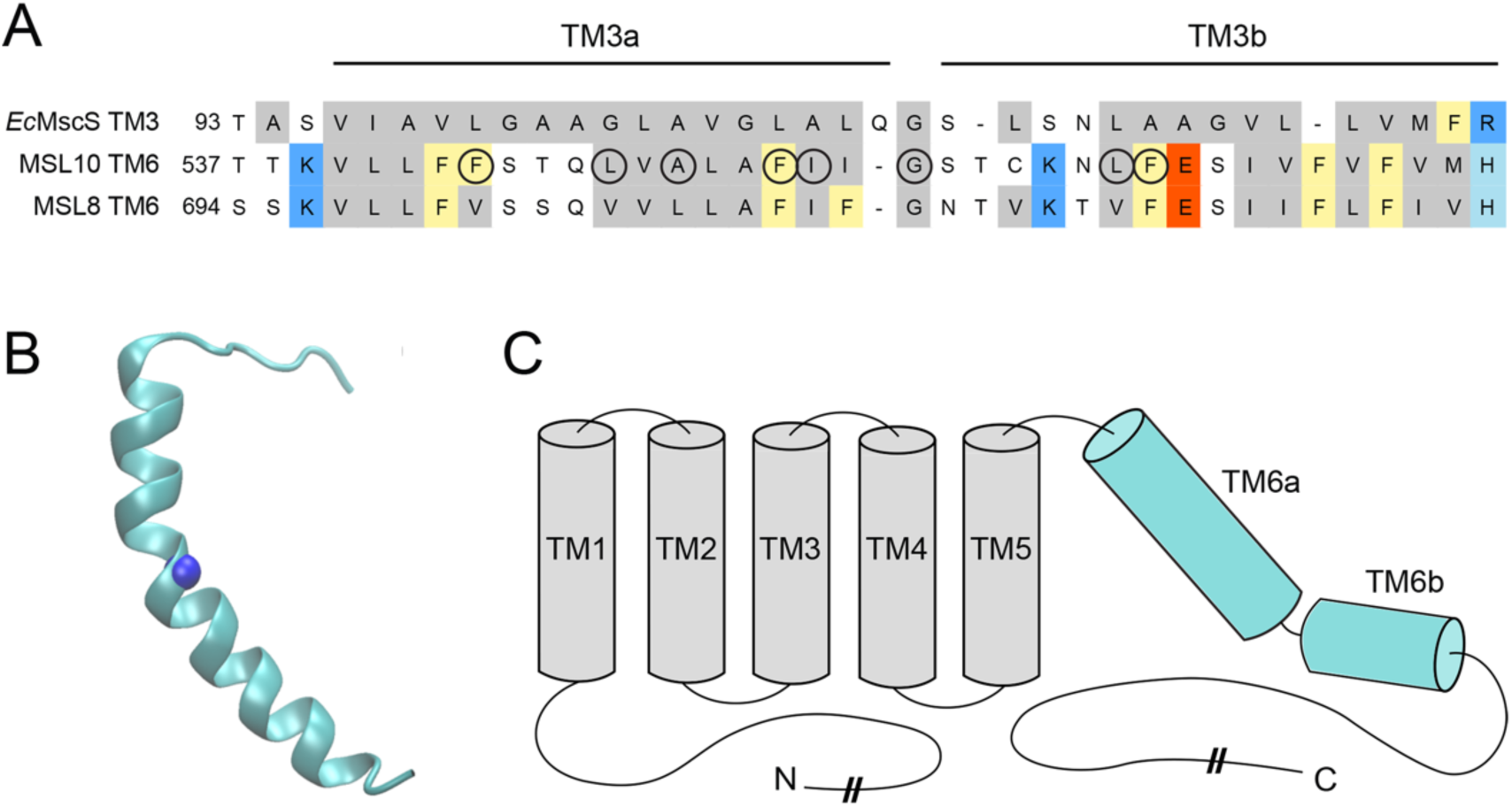
Identification of potential pore-disrupting residues in MSL10. **(A)** Alignment of the pore-lining domain of *E. coli* MscS and corresponding regions of *A. thaliana* MSL10 and MSL8. Acidic residues are indicated in red; basic residues in blue; and nonpolar in grey. Phe residues are marked yellow. Circles indicate residues that were analyzed in this report. **(B)** Side view of the predicted structure of the MSL10 TM6, created with I-TASSER. The side chain at G556, predicted to form a kink, is indicated with a blue sphere. **(C)** Predicted topology of MSL10, indicating soluble N- and C-termini and six membrane-spanning helices. The length of the N- and C-termini are not to scale.

### Phenylalanine 553 maintains channel conductance and the stability of the open state

One distinction between the amino acid sequences of *Ec*MscS TM3 and MSL10 TM6 is the presence or absence of multiple phenylalanine residues. *Ec*MscS TM3 contains only one Phe residue, and it is located at very end of TM3b. MSL10 has six Phe residues scattered through the pore-lining domain. MSL8 has six Phe residues in its TM6; five of these are conserved with MSL10 (indicated in yellow, Figure 1C), and F720 is essential for channel function ^58^. To determine if Phe residues in TM6 are also critical for MSL10 channel function, we used site-directed mutagenesis to change F544, F553, and F563 to smaller nonpolar residues. We introduced F544V, F553W, F553L, F553V, and F563L lesions into the MSL10 coding sequence of pOO2-MSL10-GFP, and pOO2-MSL10 for in vitro capped RNA (cRNA) production^53^. cRNA for all variants was injected into Xenopus oocytes for expression and characterization as previously described^43^.

To determine if these lesions affected protein folding or stability, we monitored the trafficking of MSL10 variants fused to GFP by confocal microscopy. All variants produced similar signal at the oocyte periphery 5 days after injection, while no signal was present in water-injected oocytes (Figure 2A). Single channel patch-clamp electrophysiology was then used to assess the channel behavior (voltage-dependent conductance, open state stability, and tension-sensitivity) of each of these MSL10 variants, as in ^59^. Current traces from excised inside-out patches derived from oocytes expressing wild type MSL10 showed the expected stable single channel openings (119.4 ± 4.0 pS at -100mV, Figure 2B, top). MSL10 F544V did not appear different from the WT. However, MSL10 F553W, MSL10 F553L, and MSL10 F563L exhibited “flickery” channel activity, defined here as rapid increases and decreases in conductance without a clear time spent in the open state (middle and bottom, Figure 2B). We were unable to detect any channel activity in oocytes expressing MSL10 F553V.

**Figure 2.**
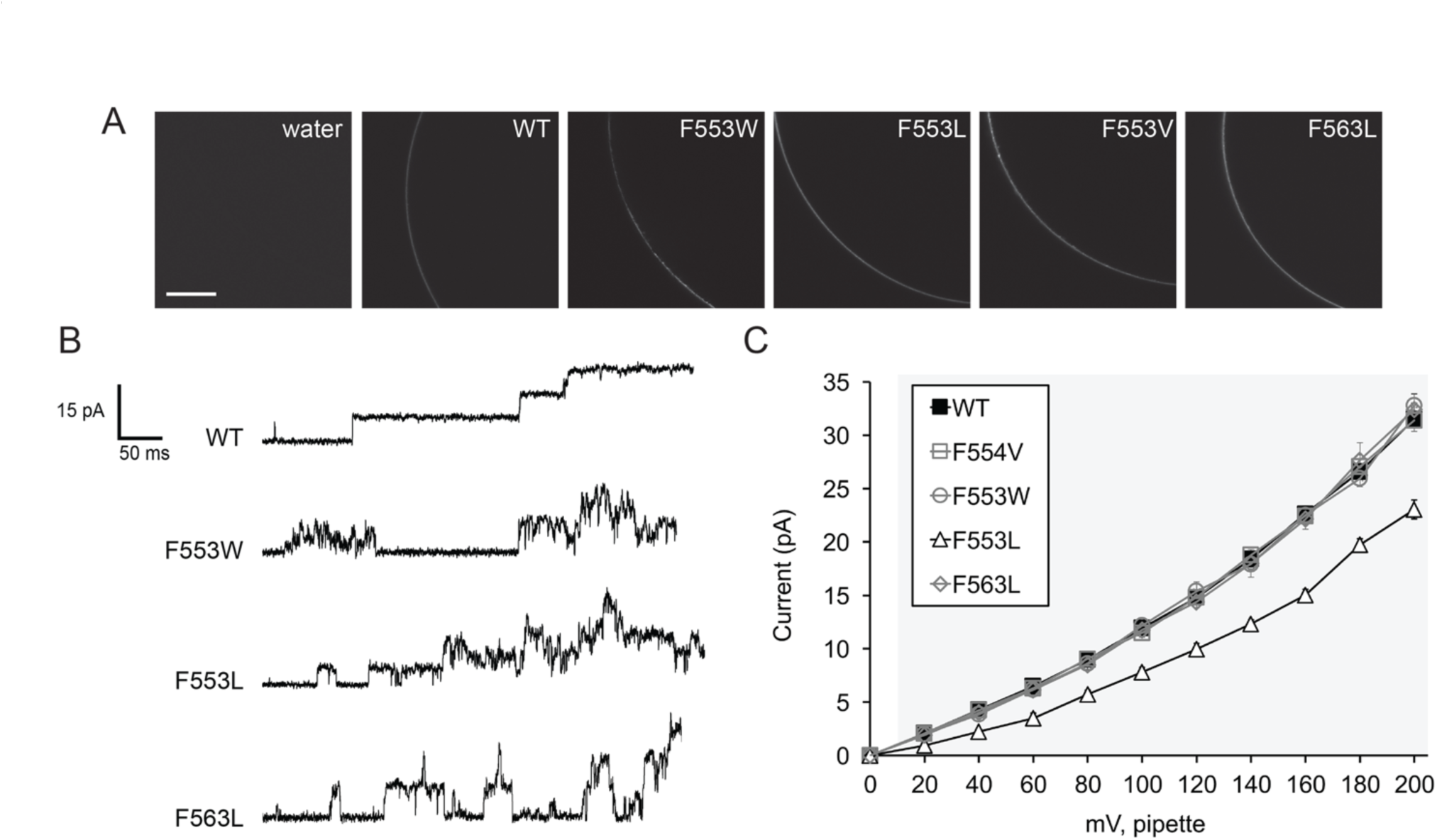
Mutagenesis of two TM6 phenylalanines reveals a role for bulky nonpolar residues in maintaining channel conductance and the stability of the open state. **(A)** Portions of the oocyte periphery 5 days after injection with the indicated MSL10-GFP variant cRNA. Scale bar is 100 μm. (**B**) Examples of traces at -100 mV membrane potential. 0.5 second fragments of 5 s records of channel activation by symmetric pressure ramps are shown. (**C**) Current/voltage curves for the indicated MSL10 variants. Each data point is the average current from 3–9 patches in 60 mM MgCl_2_, 4 mM HEPES. Error bars indicate standard deviation but are obscured by the symbols. Grey background indicates voltages where the current produced by MSL10 F553L differed significantly from wild type MSL10, p < 0.001 (Student’s t-test).

We measured the unitary currents produced by MSL10 and MSL10 Phe variant channels under transmembrane voltage from 0 to -200 mV. The unitary conductances of MSL10 F544V (114.8 ± 4.0 pS), F553W (121.8 ± 4.3 pS) and F563L (117.7 ± 5.4 pS), calculated at -100 mV membrane potential, were indistinguishable from wild type MSL10 (Figure 2C, Table 1). However, MSL10 F553L produced single-channel currents that were significantly lower than that of the wild type at every voltage tested (Figure 2C), and its conductance at -100mV was 78.0 ± 4.4 pS, 0.6-fold that of the wild type (Table 1). As noted above, MSL10 F553V did not produce any activity. Thus, successively reducing the bulkiness of residues at position 553 resulted in successively lower unitary channel conductance (W≈F >L>V). We conclude that F553 and F563 both play important roles in maintaining the stability of the open state of MSL10, while F553 also controls channel conductance.

**Table 1.**
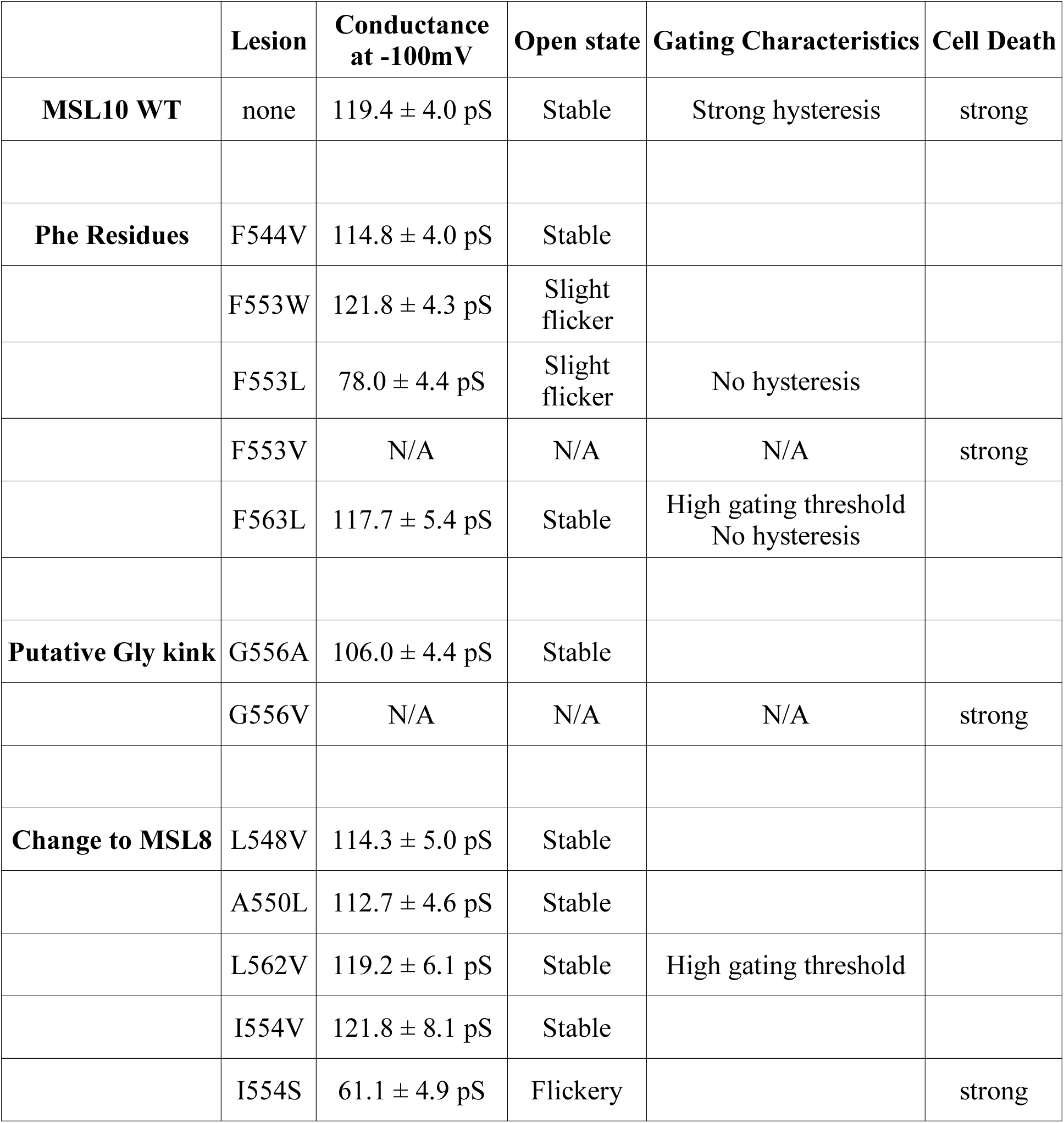
Properties of the *At*MSL10 TM6 mutants. Conductance at -100mV.

|  | **Lesion** | **Conductance**<br>**at -100mV** | **Open state** | **Gating Characteristics** | **Cell Death** |
| --- | --- | --- | --- | --- | --- |
| **MSL10 WT** | none | 119.4 ± 4.0 pS | Stable | Strong hysteresis | strong |
| **Phe Residues** | F544V | 114.8 ± 4.0 pS | Stable |  |  |
|  | F553W | 121.8 ± 4.3 pS | Slight<br>flicker |  |  |
|  | F553L | 78.0 ± 4.4 pS | Slight<br>flicker | No hysteresis |  |
|  | F553V | N/A | N/A | N/A | strong |
|  | F563L | 117.7 ± 5.4 pS | Stable | High gating threshold<br>No hysteresis |  |
| **Putative Gly kink** | G556A | 106.0 ± 4.4 pS | Stable |  |  |
|  | G556V | N/A | N/A | N/A | strong |
| **Change to MSL8** | L548V | 114.3 ± 5.0 pS | Stable |  |  |
|  | A550L | 112.7 ± 4.6 pS | Stable |  |  |
|  | L562V | 119.2 ± 6.1 pS | Stable | High gating threshold |  |
|  | I554V | 121.8 ± 8.1 pS | Stable |  |  |
|  | I554S | 61.1 ± 4.9 pS | Flickery |  | strong |

### Glycine 556 is a key residue for MSL10 channel function

The next residue in MSL10 we chose to investigate was G556. MSL10 G556 aligns with the kink-forming residue G113 in *Ec*MscS (Figure 1A) and is predicted to sit at a similar kink in TM6 (Figure 1B). To determine if G556 plays a role in MSL10 channel function, we changed this residue to the larger nonpolar residues alanine (G556A) or valine (G556V). While both MSL10 variants were expressed in Xenopus oocytes and trafficked normally to the plasma membrane (Figure 3A), neither functioned like the wild type. The MSL10 G556A mutant was active and produced a relatively stable channel opening (Figure 3B). However, it had a unitary conductance of 106.0 ± 4.4 pS, 0.9-fold that of the wild type channel (Figure 3C). MSL10-G556V did not produce any activity even under extreme membrane tensions and high membrane potentials. Thus, the G556A substitution produced a modest effect, and G556V completely ablated function, presumably because of increasing size of the side chains. These results establish that G556 plays a key role in the function of MSL10 and are consistent with the prediction that there is bending at this residue within the helix and that mobility at this site is important for the conformational changes associated with channel opening.

**Figure 3.**
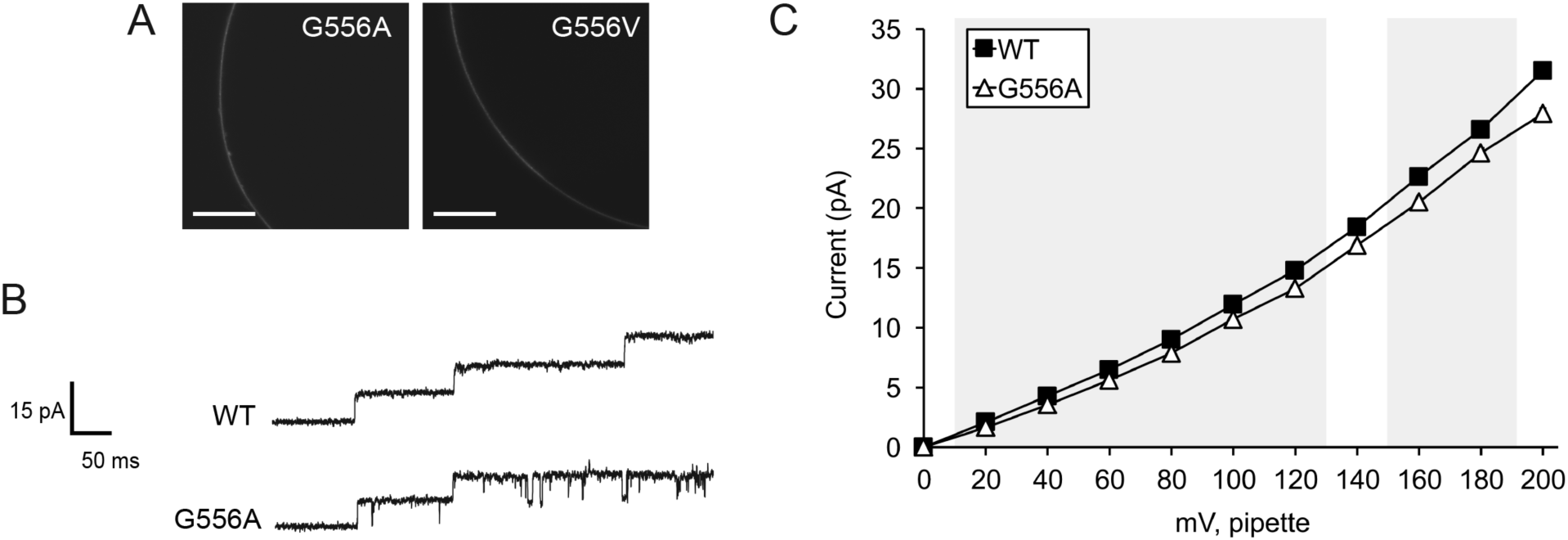
Replacing G556 with larger nonpolar residues reduces or ablates channel conductance. **(A)** Portions of the oocyte periphery 5 days after injection with the indicated MSL10-GFP variant cRNA. Scale bar is 100 μm. (**B**) Examples of traces at -100 mV membrane potential. 0.5 second fragments of 5 s records of channel activation by symmetric pressure ramps are shown. (**C**) Current/voltage curves for wild type and MSL10 G556A. Each data point is the average current from 3–9 patches in 60 mM MgCl_2_, 4 mM HEPES. Error bars indicating standard deviation are present but are obscured by the symbols. Wild type MSL10 curve is from the same data as in Figure 2C. Grey background indicates voltages where the current produced by MSL10 G556A differed significantly from wild type MSL10, p < 0.001 (Student’s t-test).

### Nonpolar TM6 residues that differ between MSL10 and MSL8 are not required for wild-type conductance and open state stability

Nonpolar residues in MSL10 TM6 that are not conserved in MSL8 include F544, L548, A550, I555 and L562. To determine if these residues confer the differences in conductance and tension sensitivity between MSL10 and MSL8, we changed several of these residues in MSL10 to the corresponding residue found in MSL8. MSL10 F544V was analyzed with other Phe substitutions in Figure 2 and did not appreciably affect MSL10 channel behavior. MSL10 L548V (114.3 ± 5.0 pS), MSL10 A550L (112.7 ± 4.6 pS) and MSL10 L562V (119.2 ± 6.1 pS) were also indistinguishable from the wild type (119.4 ± 4.0 pS) with respect to unitary conductance and open state stability (Figure 4 A and B, Table 1).

**Figure 4.**
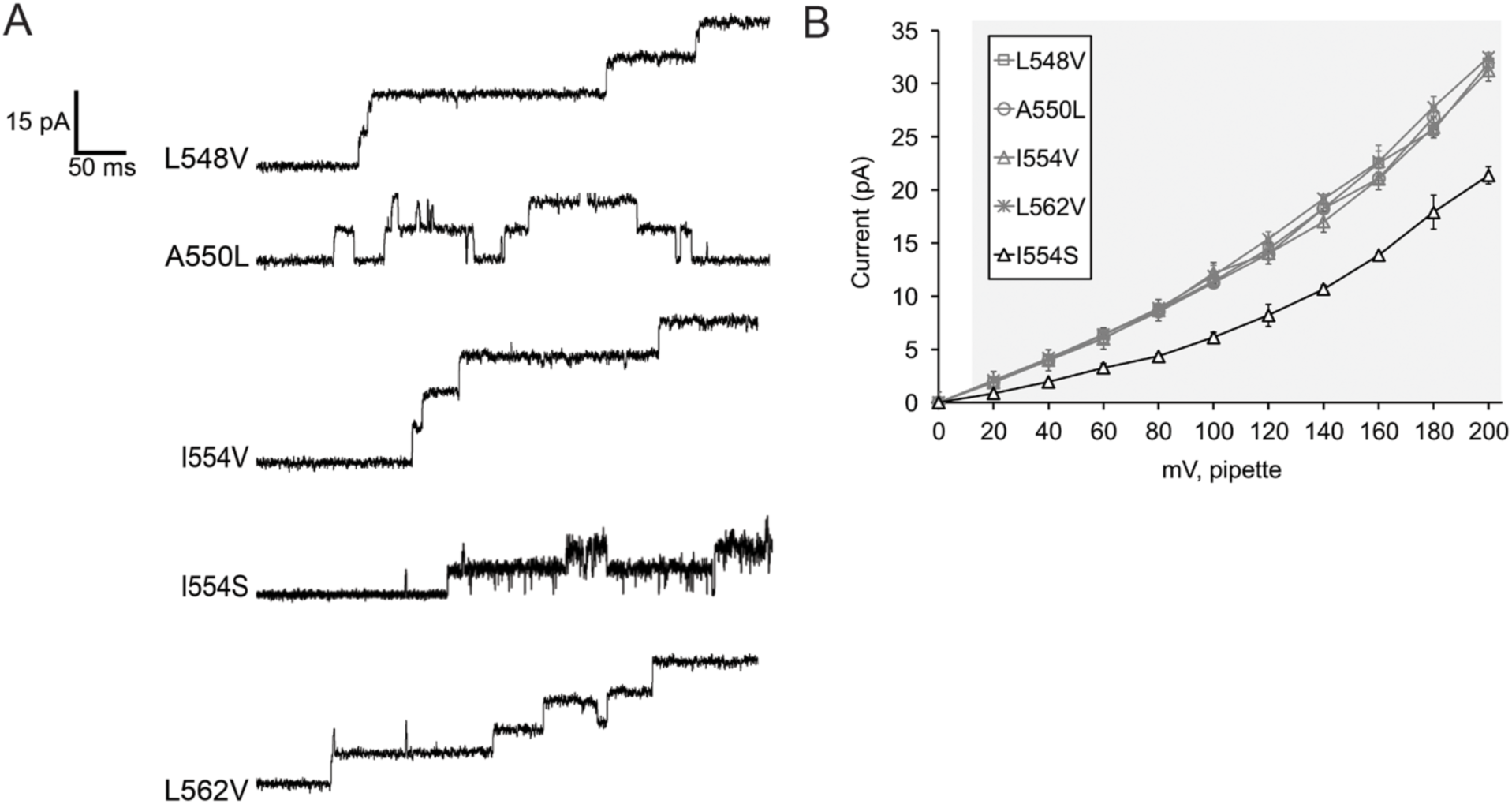
Effects of Mutating Nonpolar Residues in TM6 that Differ Between MSL8 and MSL10. (**A**) Examples of traces at -100 mV membrane potential. 0.5 second fragments of 5 s records of channel activation by symmetric pressure ramps are shown. (**B**) Current/voltage curves for the indicated MSL10 variants. Each data point is the average current from 3–9 patches in 60 mM MgCl_2_, 4 mM HEPES. Error bars indicate standard deviation. Grey background indicates voltages where the current produced by MSL10 I554S differed significantly from wild type MSL10, p < 0.001 (Student’s t-test).

### Changing Isoleucine 554 to Serine disrupts channel conductance and the stability of the open state

We previously showed that MSL8 I711S produced a channel with wild type conductance but an increased tension threshold ^59^. We mutated the corresponding residue in MSL10, I554, to valine and serine. While MSL10 I554V exhibited a wild type unitary conductance (121.8 ± 8.1 pS) and gating pattern, MSL10 I554S was very flickery (Figure 4A), and had a significantly lower unitary conductance than WT (61.1 ± 4.9 pS compared to 119.4 ± 4.0 pS of WT at -100 mV membrane potential, Figure 4 B). These unexpected results suggest that MSL10 I554S has normal tension sensitivity, but once open, has an unstable open state. Taken together, the data shown in Figure 4 suggest that the differences in MSL8 and MSL10 channel behavior cannot be easily attributed to individual amino acid residues within TM6.

### Opening and/or closing tension sensitivities were altered in MSL10 F553L, L562V, F563L

The established approach for measuring tension sensitivity of *Ec*MscS, by calculating the activation midpoint (e.g. ^53^), is not possible with MSL10 expressed in oocytes, as one typically cannot reach current saturation before the patch collapses ^28,52,60^. Instead we measured the gating threshold, or the amount of tension required to open the second channel in the patch. We used pipettes with the same resistance (3.00 ± 0.25 MOhm) to reduce variability in patch size and geometry.

We first analyzed the gating threshold for wild type MSL10. We observed that the gating threshold depended on the number of channels we observed in a patch prior to saturation or patch rupture, and that patches with more channels had a lower opening threshold (Figure 5A). Assuming low open probability, and minimal spontaneous gating at zero tension, these data can be fit to a line with slope 0.049 ± 0.02 channels/mm Hg (Figure 5A, circles). The same analyses for MSL10 F553L (squares) and MSL10 L562V (triangles) produced slopes similar to that of the wild type channel, 0.047 ± 0.04 and 0.036 ± 0.01 channels/mm Hg respectively (Figure 5A). However, MSL10 F563L did not show a strong dependence of channel number on threshold pressure, with patches containing a range of channel numbers all opening at essentially the same tension. The linear regression line slope for MSL10 F563L was only 0.009 ± 0.007 channels/mm Hg (Figure 5B, diamonds). As shown in Figure 5B, we routinely observed very high channel numbers per patch for MSL10 F563L in some cases as many as 600 channels per patch. Thus, some aspect of channel-channel interaction may be altered in MSL10 F563L.

**Figure 5.**
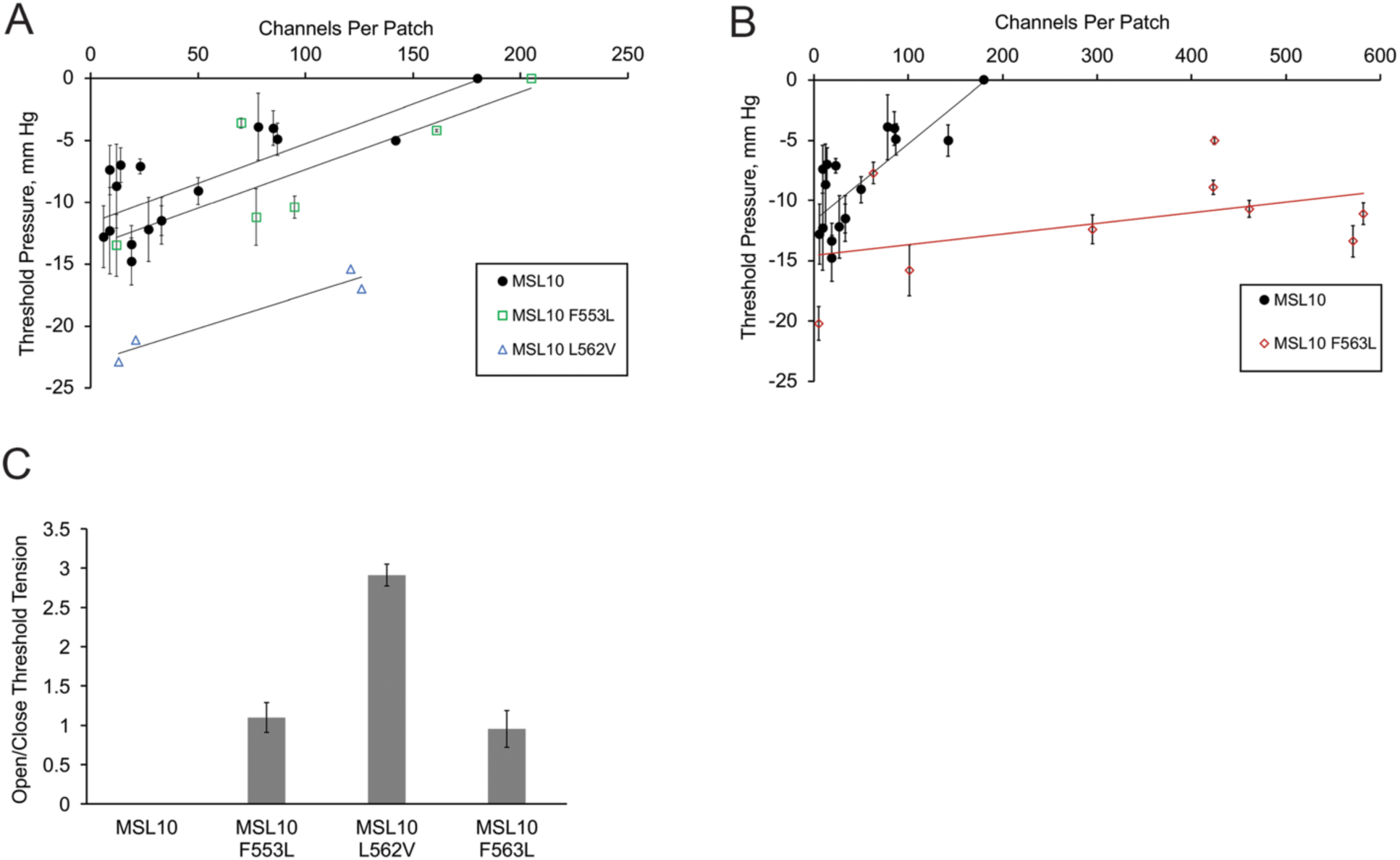
Three Bulky Nonpolar Residues Affect the Threshold Tensions of Opening and Closing (A, B) Plot of channel number versus threshold tension for wild type MSL10 and the indicated variants. Data were corrected for access resistance. The data for wild type MSL10 are the same in both plots. Error bars indicate standard deviation between the threshold pressure measured from 5–10 trials from a single patch containing the indicated number of channels. **(C)** The average open/close membrane tension ratios for MSL10 and variants including data from all channel number patches shown in panels A and B. One sample for MSL10 F553L was left out as outlier. Error bars indicate standard deviation between the average open/close ratio for each patch.

The fact that all three MSL10 variants had higher closing thresholds than wild type MSL10 made it possible to calculate an open/close ratio, using patches with 5-600 channels (Figure 5C). For MSL10 F563L, the open/close ratio was close to one (0.95 ±0.23). MSL10 L562V had an open/close ratio of 2.9 ± 0.13. For MSL10 F553L, the open/close ratio was close to one (1.10 ± 0.19) when only the 4 closest data points were considered. A fifth strongly outlying data point was not included, as its value exceeded the upper fence, as defined by the quartile method. To summarize, wild type MSL10 showed threshold pressure of around -15 mm Hg and exhibited hysteresis. MSL10 L562V had a high tension threshold for opening but maintained hysteresis. MSL10 F563L had a high tension threshold for both opening and closing and did not exhibit hysteresis. Due to the small sample size, we can only tentatively conclude that MSL10 F553L lost hysteresis.

### MSL10 TM6 lesions that ablate channel function do not alter the ability of MSL10 to trigger cell death signaling

These lesions that alter MSL10 channel behavior provide tools to test the structural requirements for MSL10’s cell death signaling function. The two MSL10 lesions that produced no ion channel activity (F553V and G556V), along with wild type MSL10, were fused to GFP and transiently expressed under the strong constitutive Cauliflower Mosaic Virus 35S promoter (35Sp) in tobacco leaf epidermal cells as described previously ^43^. Constructs were co-infiltrated with a plasmid expressing P19 in order to suppress gene silencing ^54^. The As shown in Figure 6A, all MSL10 variants were expressed and trafficked normally in tobacco cells. Five days after Agrobacterium infiltration, leaf samples were dual stained for FDA and PI to assess cell death (Figure 6B). As expected, 35% of the cells in leaf samples expressing wild type full-length MSL10 were dead, compared to 14% when P19 was infiltrated alone. Neither MSL10-F553V nor MSL10 G556V produced levels of cell death that were statistically distinguishable from wild type MSL10 (Figure 6C), indicating that ion channel function is not required for the ability of full length MSL10 to trigger cell death in this transient assay.

**Figure 6.**
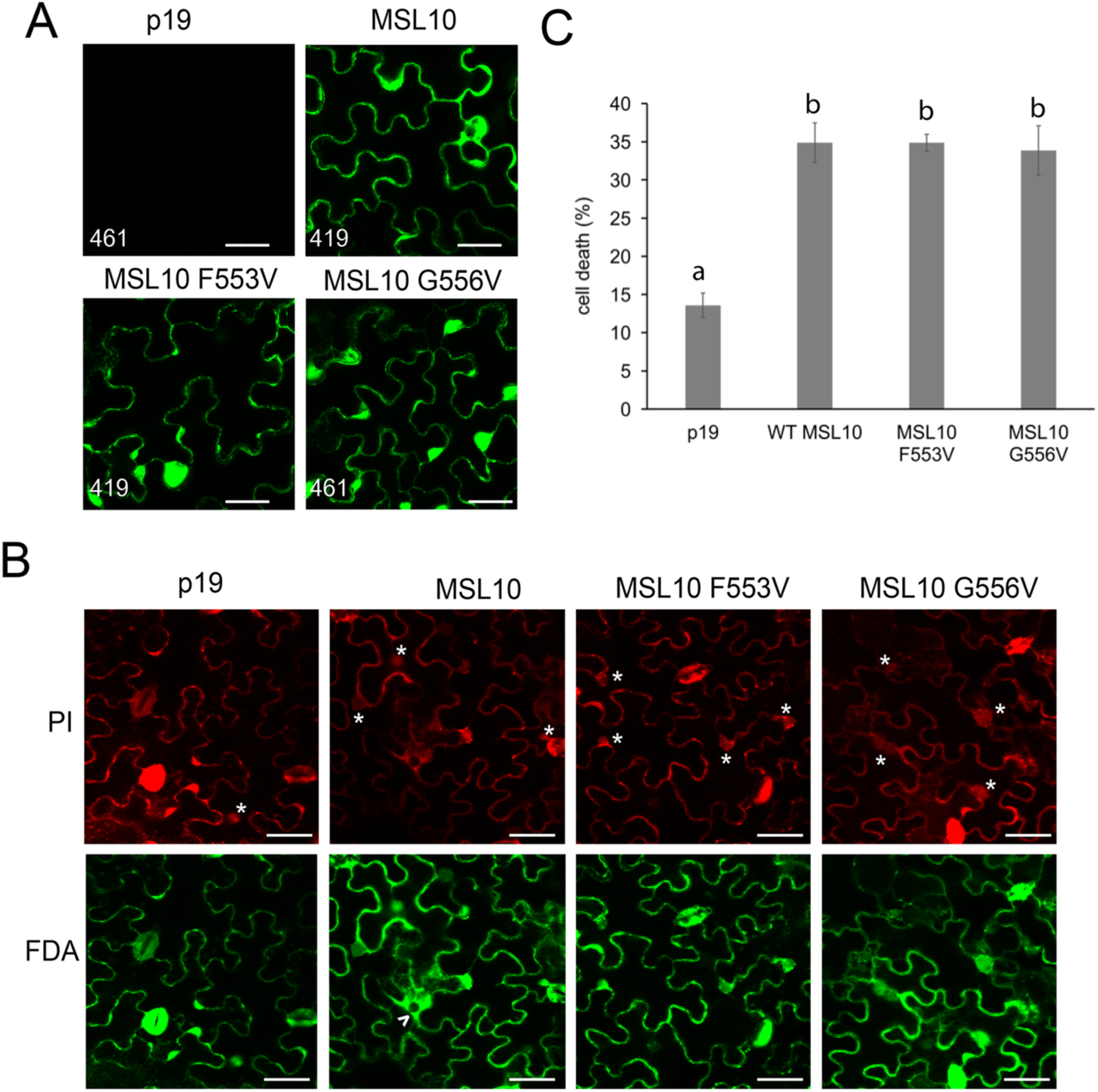
MSL10 Mechanosensitive channel activity is not required for death signaling in planta. **(A)** Localization of MSL10-GFP variants transiently expressed in tobacco epidermal pavement cells. Images were taken five days after infiltration. Infiltrations with P19 alone were used as a negative control. Scale bar is 40 μm. Numbers in the left-hand corner indicate the voltage setting of the PMT detector when imaging. **(B)** Examples of cell viability assays. Tobacco leaves expressing MSL10-GFP variants were dual stained with FDA and PI; cells were scored as dead if they had a PI-stained nucleus (indicated by asterisks) or a disappearing vacuole, evidenced by spreading of cytoplasmic signal (arrowhead). Scale bar is 50 μm. **(C)** Percentage of dead cells quantified from dual staining of 20 leaves (P19 and MSL10) or 10 leaves (MSL10 variants) from multiple infiltration experiments. Error bars indicate standard error. Statistical differences were assessed by One-way ANOVA and Scheffe’s test; groups with the same letter (b) did not significantly differ from each other (p>0.05).

## Discussion

Here we report the functional effect of twelve different point mutations in the mechanosensitive channel MscS-Like (MSL)10 from *Arabidopsis thaliana*. All substitutions were made in a set of eight non-polar amino acids located in the putative pore-lining transmembrane (TM) helix 6. We observed that four of these lesions (F544V, L548V, A550L, and I554V) did not detectably alter channel behavior in our assays, four primarily affected unitary conductance and/or open state stability (F553W, F553L, G556A, I554S), three affected tension sensitivity and/or hysteresis (F553L, L562V, and F563L), and two produced no mechanosensitive channel activity at all (F553V, G556V). Finally, we observed that the latter two variants were as capable as wild type MSL10 of inducing cell death in an *in planta* expression assay.

Since Phe residues are found in TM6 of MSL8 and MSL10 but not in TM3 of *Ec*MscS, they were reasonable candidates for residues that underlie the differing channel properties of MSLs and *Ec*MscS. MSL10 has a smaller conductance than *Ec*MscS, and we originally hypothesized that these bigger hydrophobic residues found in the MSL10 TM6 could form a “vapor lock”, like *Ec*MscS L105 and L109 ^22,61^, or partially obstruct the channel pore as suggested for F450 in the Arabidopsis SLAC1 channel ^62^. In this case, replacing Phe with smaller side chains such as Leu or Val would lead to a deregulated or constitutively open channel. Alternatively, bulky Phe residues could be involved in the slow closing kinetics observed in MSL10 but not in MscS if they participated in a hydrophobic residue-mediated force transduction pathway from the periphery of the channel to its pore, in a manner analogous to the proposed force-transmitting clutch of *Ec*MscS ^32^. However, none of these hypotheses were correct. Instead, our data suggest that the large hydrophobic nature of the Phe residues at 553 and 563 facilitate the unitary conductance and the stability of the open state of MSL10.

While mutation of F553 to tryptophan had no significant effect on conductance, mutation to leucine led to a decrease in unitary conductance and mutation to valine produced a channel without any apparent mechanosensitive response (Figure 2). Because MSL10 F553L produced a partially functional channel, and because MSL10 F553V was expressed at wild-type levels, MSL10 F553V is likely to produce a stable oligomeric channel that lacks conductive pore. We thus speculate that the pore size of MSL10 is altered by the size of the side chain at F553; as the side chains at F553 decrease in size, pore-lining domains grow more and more closely packed, thus successively reducing conductance.

While gating pressures for the Phe mutants did not significantly differ from those of WT, both MSL10 F553L and MSL10 F563L showed almost no hysteresis (Figure 5C). MSL10 typically shows delayed closing relative to opening kinetics; with MSL10 F553L and MSL10 F563L, the ratio of opening and closing tensions were close to 1. This suggested that these mutant channels have an open state that transitions back to the closed state more easily than the wild type channel. The flickery behavior of MSL10 F553L may be explained by a steric mismatch between the mutated residue and its interaction partner from the adjacent pore-lining domain. Lack of any channel activity for the F553V mutant may indicate insufficient size of the non-polar side chain for creating even partially open pore. We speculate that the open state is stabilized by interaction between F553 and another hydrophobic residue from the adjacent pore-lining domain of the channel. Taken together, the data shown in Figure 2 show that multiple Phe residues in MSL10 TM6 are required for the size and stability of the open pore.

We also found that G556 plays a critical role in MSL10 channel function. We first targeted G556 for mutation because glycine at a similar position in *Ec*MscS TM3 (G113) has been implicated in gating movements through crystal structures and molecular dynamics. The *Ec*MscS G113A lesion prevents channel inactivation, presumably by preventing the helix kinking thought to accompany MscS entry from the open state into the inactivated state ^16,17^. While MSL10 does not show inactivation in oocytes (though it does in plant cells ^63^), the model for MSL10 TM6 shown in Figure 1 suggests a similar geometry and is consistent with a similar gating movement in the pore-lining helix between MscS and MSL10. We therefore hypothesized that lesions at this site might affect the conformational changes required for channel gating.

While MSL10 G556A showed only a subtle difference from the wild type—a small, but statistically significant, decrease in conductance—MSL10 G556V did not produce any mechanosensitive activity at any applied tensions, though the mutant was expressed at WT levels (Figure 3). We speculate here that the MSL10 G556 side chain faces the pore lumen in the open state and therefore introduction of alanine at position 556 resulted in decrease of unitary currents (Figures 3B, C). Introduction of the even bigger hydrophobic residue valine at this site might then either fully occlude the channel pore, or prevent the conformational changes associated with channel gating. Whether the same movements are made during the gating transition of *Ec*MscS and MSL10 is not clear and will certainly require additional experimentation, but the fact that the G556 is a key structural feature of MSL10 TM6 is established.

We addressed the similarities and differences between MSL10 and its close homolog MSL8. We first attempted to link the difference in MSL8 and MSL10 channel characteristics to the differences in the sequence of their putative pore-lining domains. We mutated 3 residues of MSL10 into their counterpart in MSL8, generating MSL10 L548V, A550L, and L562V. Only the latter had any effect; MSL10 L562V had a high gating threshold (Figure 5A), similar to that previously observed with MSL8 ^44^. Thus, L562/V719 may in part be responsible for the difference between MSL8 and MSL10 with respect to tension sensitivity.

We also made mutations in residues conserved between MSL8 and MSL10 that have known effects on MSL8 function. MSL8 I711S exhibited normal conductance but an increased gating threshold and MSL8 F720L was a completely disrupted channel ^53^. Both lesions altered MSL10 function, but they did not produce the same effects they did in MSL8. MSL10 I554S (aligns with MSL8 I711) produced a flickery channel with half the conductance of the wild type, while MSL10 F563L (aligns with MSL8 F720) produced a channel with normal conductance but a high gating threshold. These unexpected results suggest that local or global differences in structure dictate the characteristics of these two channels, so trying to identify individual residues responsible for particular channel characteristics may not be successful.

Residues L562 and F563 form a hydrophobic patch flanked by polar and charged residues (Figure 1A) and our experiments indicate that they are essential for normal tension sensitivity and gating kinetics of the MSL10 channel (Figure 5). Hysteresis is the strongest in WT MSL10, wherein the threshold tension for closing is much lower than for opening. The difference between opening and closing tensions was decreased in MSL10 F562V, and completely abolished in MSL10 F563L (Figure 5C). We speculate that these two residues may function as a “force transmitters”, similar to L111 of *Ec*MscS TM3. It has been proposed that L111 interacts with F68 from TM2, enabling transduction of the membrane tension into a gating force ^32^. Similarly, L562 and F563 residues may be part of an intra-transmembrane helix system that serves to control the closing kinetics of MSL10 by stabilizing the open state.

The concept that ion channels have functions separable from their ability to mediate ion flux (non-conducting functions) is not new, and has been established for sodium and potassium channels in animal systems ^64,65^. However, non-conducting functions have not been previously demonstrated for MS ion channels. We previously established that MSL10 is capable of inducing cell death when overexpressed in stable Arabidopsis lines or in transient tobacco expression experiments ^54^. This effect was modulated by seven phosphorylation sites in the soluble N-terminus, but these sites did not alter ion channel function, suggesting that the two functions might be separable. Transient overexpression of the soluble N-terminus of MSL10 was capable triggering programmed cell death on its own. However, these results left open the possibility that MSL10 ion channel function is required indirectly in order to activate the cell death signaling mediated by the N-terminus, and that this effect could be simulated by expressing the truncated, soluble N-terminus.

Two of the twelve lesions we tested, MSL10 F553V and MSL10 G556V, were normally expressed and trafficked in Xenopus oocytes but did not exhibit any mechanosensitive ion channel activity (Figures 2 and 3). These mutants provided the opportunity to test the idea of a non-conducting function for MSL10 more directly and without the caveats of the truncation experiment. The effect of overexpressing these mutant channels did not differ in any way from the wild type, indicating that MS ion channel activity is not required for MSL10’s ability to induce cell death in tobacco epidermal cells. The results presented here now firmly establish that MSL10 induces programmed cell death through a mechanism independent of mechanically-induced ion flux. A key future experiment will be to determine if the non-conducting function of MSL10 is regulated by membrane tension. It will also worth testing if non-conducting functions are a conserved feature of proteins in the MscS family, as suggested by a report that the soluble C-terminus of MscS interacts with the bacterial fission protein FtsZ ^66^.

## Methods

### Molecular biology

All constructs were based on pOO2-MSL10 ^43^. Site-directed mutagenesis was used to introduce point mutations into the *MSL10* sequence and confirmed by sequencing. Capped cRNA was transcribed *in vitro* with SP6 polymerase using the mMessenger mMachine kit (Ambion, Thermo Fisher Scientific) and stored at -80°C at approximately 1000 ng/μl.

### Oocyte preparation

*Xenopus laevis* oocytes (Dumont stage V or VI) were purchased (Ecocyte Bioscience US LLC, Austin, Texas) and handled as described ^59^. Oocytes were injected with 50 nl of 1000 ng/μl of RNA the day after isolation. Fluorescent imaging of the oocytes was carried out 48–72 hours after injection. Briefly, the oocytes were placed on concaved slides and covered with coverslips. Confocal imaging of the periphery of the oocytes was performed using Olympus Fluoview 1000 with BX61 microscope and the Olympus FV10-ASW software suite.

### Electrophysiology

The buffer used was 60 mM MgCl_2_, 5 mM HEPES, adjusted to pH 7.38 with TEA-OH. All the traces presented were obtained from excised inside-out patches. Data were acquired using Axopatch 200B amplifier and Digidata 1440A digitizer (Molecular Devices) at 20 kHz and low-pass filtered at 5 kHz. Channels were activated by symmetric 5-second pressure ramps. Pressure was applied and monitored with a HSPC-1 high speed pressure clamp system (ALA Scientific Instruments), and traces analyzed with Clampfit 10.6 (Molecular Devices). The gating threshold of a channel variant was defined as the pressure at which the second channel of the population in a patch opened. For each patch, several pressure ramps of -30 mmHg were run to accommodate for patch creep. Only after that, measurements at -20 mV membrane were performed. For each patch the results of 7–12 consecutive pulls were averaged. The number of channels per patch was estimated from the peak current at patch rupturing pressure. In cases when the number of open channels was 100 or more, the correction for series pipette resistance was introduced. For closing pressures, the average was taken only in cases when closing pressure was not zero. In case when at least one pull for a patch resulted in at least one open channel after the pressure was released, the closing pressure was considered to be zero.

### Software

The putative structure of *At*MSL10 TM6 (Fig. 1B) was obtained from the I-TASSER prediction server ^56^ using the *Ec*MscS closed state crystal structure (2OAU:A, ^22,57^) as a template. Sequence alignments and analysis were made using Unipro UGENE bioinformatics toolkit ^67^. Visualization of crystal structures and imaging of the putative *At*MSL10 pore region were performed in VMD suite ^68^. Secondary structure of *At*MSL10 was predicted using ARAMEMNON plant membrane protein database ^58^.

### Cell death assays

The coding sequences of wild type MSL10, MSL10 F553V, and MSL10 G556V were amplified from the pOO2 vectors describe above and cloned into the pK7FWG2 vector for C-terminal GFP tagging and transient expression in *Nicotiana benthamiana* leaves under the control of the 35S promoter. In order to quantify the amount of dead versus viable epidermal pavement cells, leaves were dual stained with fluorescein diacetate (FDA) and propidium iodide (PI) five days after infiltration as described in ^69^ and visualized via confocal microscopy. As previously outlined, cells were considered dead 1) if their nucleus was stained by PI and/or 2) their vacuole disappeared, which is apparent when cytoplasmic FDA/GFP signal fills the entire body of the pavement cell ^54^. In this study, we added a third criterion whereby cells were also considered dead if their vacuole had significantly, though not completely, disappeared. The % cell death reported was the average of four separate infiltration experiments, each consisting of three to four leaves per construct, with n ∼ 60 cells imaged per leaf.

## Author Contributions

GM, JMS and ESH designed experiments and wrote the paper; GM, JMS and SO performed experiments.

## Acknowledgments

We acknowledge Debarati Basu for generating the mutant MSL10 constructs used in Figure 6. These experiments were funded by the National Institutes of Health grant R01GM084211 (to ESH), National Science Foundation grant MCB1253103 (to ESH) and NSF Graduate Research Fellowship DGE-1745038 (to JMS).

